# microRNA-184 and Its lncRNA Sponge UCA1 are Induced in Wounded Keratinocytes in a Store-Operated Calcium Entry-Dependent Manner

**DOI:** 10.1101/328401

**Authors:** Adam Richardson, Daniel Owens, Kehinde Ross

**Affiliations:** School of Pharmacy and Biomolecular Sciences, Liverpool John Moores University, Liverpool, United Kingdom.; School of Sport and Exercise Sciences, Liverpool John Moores University, Liverpool, United Kingdom.

**Author notes:** **Correspondence:** Kehinde Ross, School of Pharmacy and Biomolecular Sciences, Liverpool John Moores University, Liverpool, L3 3AF, England.

**Keywords:** MicroRNA, store-operated calcium entry, long non-coding RNA, UCA1, cell migration

## Abstract

Emerging evidence implicates microRNAs (miRNA) in the regulation of keratinocyte migration. However, the putative roles of microRNA-184 (miR-184) in keratinocyte migration have not been examined. Here, we show that miR-184 expression was elevated following wounding of human keratinocyte monolayers. The induction of miR-184 was dependent on store-operated calcium entry (SOCE) as it was abolished by pharmacologic SOCE blockers. The long non-coding RNA urothelial cancer associated 1 (UCA1), which is thought to acts as a sponge or competing endogenous RNA (ceRNA) against miR-184 was also induced in scratched monolayers. Induction of UCA1 was impaired, but not abolished, by SOCE inhibition. Transfection of keratinocytes with a miR-184 mimic stimulated migration in scratch assays, whereas inhibition of miR-184 dampened the ability of keratinocytes to migrate. Together, our data suggest, for the first time, that SOCE promotes miR-184 induction in wounded monolayers to support keratinocyte migration while also increasing lncRNA UCA1 expression, which may in turn regulate miR-184 activity in keratinocytes.

## INTRODUCTION

Despite significant advances in understanding the multiple factors associated with chronic wounds, there remains a significant unmet need for therapeutic interventions to promote wound healing. With the ageing population and associated rise in the incidence of diabetes, pressure ulcers, venous leg ulcers and diabetic foot ulcers, the clinical and socioeconomic challenges presented by non-healing wounds are likely to persist, exerting enormous pressure on health services in both industrialised and developing nations (Eming et al., 2014, Nunan et al., 2014, Whittam et al., 2016). Estimates put the costs of managing wounds and associated comorbidities at £5.3 billion annually in the UK (Guest et al., 2015) and a staggering $25 billion in the USA (Sen et al., 2009).

The formation of new epidermal tissue over the denuded wound surface is fundamental to the completion of wound healing. Such re-epithelialization is mediated by keratinocyte migration from the wound edge to repopulate the exposed extracellular matrix, a processes that is impaired in chronic wounds (Usui et al., 2008). Multiple growth factors stimulate keratinocyte migration but the roles of non-coding RNAs have only recently gained attention.

MicroRNAs (miRNAs) are short ≈22 nucleotide non-coding RNA molecules that regulate gene expression in a posttranscriptional manner through mechanisms that, parsimoniously, culminate in the degradation of mRNA targets (Eichhorn et al., 2014). Several microRNAs (miRNAs) have been implicated in the regulation of keratinocyte migration, as reviewed recently (Ross, 2018). For instance. miR-21, miR-31 and miR-132 promote keratinocyte migration (Li et al., 2015, Li et al., 2017, Wang et al., 2012, Yang et al., 2011). In contrast, over expression of miR-24 or miR-483-3p inhibited HPEK migration (Amelio et al., 2012, Bertero et al., 2011). Keratinocyte migration is also regulated by miR-205, but both enhancement and inhibition of miR-205 have been linked to re-epithelization (Wang et al., 2016, Yu et al., 2010).

Long non-coding RNAs (lncRNAs) are a large, heterogeneous group of ncRNAs ≥ 200 nucleotides long with diverse functions and mechanisms of action (Dykes and Emanueli, 2017). One theory that is gaining traction is that lncRNAs can function as competing endogenous RNAs (ceRNAs) to sequester miRNAs and limit their function (Yang et al., 2016). Although some aspects of the ceRNA hypothesis have been challenge (Denzler et al., 2014), the long non-coding RNA UCA1 (urothelial carcinoma associated 1) appears to function as a ceRNA for miR-184, possessing as it does 4 miR-184-binding sites (Zhou et al., 2017).

Depletion of cellular calcium (Ca^2+^) stores from the endoplasmic reticulum (ER) is coupled to the influx of Ca^2+^ from the extracellular milieu. This process, known as store-operated Ca^2+^ entry (SOCE), is mediated by STIM1, which senses the drop in ER Ca^2+^ levels and forms clusters that activate ORAI1, the predominant SOCE channel (Hogan and Rao, 2015) (Gudlur and Hogan, 2017). Interestingly, ORAI1 is required for optimal keratinocyte migration, (Vandenberghe et al., 2013) but the relationships between SOCE and miRNAs involved in keratinocyte migration have received little attention.

We previously reported that little or no miR-184 was detected in human primary epidermal keratinocytes (HPEK) maintained as monolayer cultures in low Ca^2+^ media (Roberts et al., 2013). In contrast, we have recently observed miR-184 induction in HPEK exposed to elevated extracellular Ca^2+^ but not other differentiation agents (Richardson et al., 2018). The Ca^2+^-dependent induction of miR-184 was impaired when SOCE was blocked with pharmacologic inhibitors or with short-interfering RNA (siRNA) against ORAI1. Given that keratinocyte migration from the basal layer to the stratum corneum is a fundamental element of epidermal differentiation, we hypothesized that induction of miR-184 may contribute to keratinocyte migration and that induction of miR-184 may be accompanied by upregulation of its putative molecular sponge lncRNA UCA1.

## METHODS

### Reagents

Oligonucleotides (miR-184 mimic and non-targeting control) were obtained from GE Healthcare (Little Chalfont, UK). The locked nucleic acid (LNA) miR-184 inhibitor and a non-targeting control were purchased from Exiqon (Vedbaek, Denmark). Gadolinium(III) chloride (Gd^3+^) was supplied by Bio-Techne (Abingdon, UK). BTP2 (also known as YM58483 was purchased from Abcam (Cambridge, UK). Human progenitor epidermal keratinocytes (HPEK) were and CnT-Prime culture medium were from CellnTec (Bern, Switzerland).

### Semi-quantitative Reverse Transcriptase PCR (sqRT-PCR)

Cells were seeded at 3 × 10^5^/well of a 6-well plate and allowed to reach confluence. Monolayers were scratched with a pipette tip and maintained in standard CnT-Prime medium with low (0.07 mM) Ca^2+^ or in CnT-Prime with 1.5 mM Ca^2+^. Cells were harvested 5 days later and total RNA was isolated using AllPrep DNA/RNA/miRNA Universal reagents (Qiagen, Manchester, UK). RNA concentration was determined using a NanoDrop^™^ 2000c. Complementary DNA (cDNA) was synthesised from 400 ng of RNA using the miScript II RT kit (Qiagen) with HiFlex buffer. PCR amplification was performed with Quantifast SYBR Green and QuantiTect miRNA/universal primers from Qiagen. The primers for lncRNA UCA1 were purchased from Primer Design (Southampton, UK). All thermocycling was performed on a Rotor-Gene^®^ as follows: 95°C for 15 min, followed by 40 cycles of 94°C for 15 s, 55°C for 30 s and 70°C for 30 s. Relative expression was determined using the 2^−ΔΔCT^ relative quantification method (Livak and Schmittgen, 2001).

### Oligonucleotide Nucleofection and Time-lapse Imaging

Keratinocytes (5 × 10^5^ cells) resuspended in 100 μl of solution P3 Primary Cell nucleofection reagents (Lonza, Castleford, UK) with 100 nM of oligonucleotide were transferred to nucleofection cuvettes and pulsed on programme DS-138 of 4D-Nucleofector (Lonza, Castleford, UK). After incubation in pre-equilibrated CnT-Prime at room temperature for 10 min, the cells were transferred to 6-well cell culture plates and incubated at 37°C, 5% CO_2_ with the media replacement the following day. Monolayers were scratched with a pipette tip upon approaching confluence and monitored at 30 min intervals by time-lapse microscopy on a Leica DMI6000B microscope with a 37°C, 5% CO_2_ environmental chamber. Linear regression was performed using Prism 5.0 (GraphPad Software, CA, USA).

### Statistical Analysis

The 2^−ΔΔt^ relative quantification method should give a fold change of 1 for the calibrator used in sqRT-PCR data analysis (Livak and Schmittgen, 2001). This is always the case for a sample, but when independent triplicates from primary cells obtained from multiple donors are averaged, the fold changes may not be near 1. Although this may reflect a high degree of experimental variation (Livak and Schmittgen, 2001) it could also reflect inherent biological variation. For simplicity, and in line with increasing calls for inferential statistics to be abandoned (McShane et al., 2017, Trafimow et al., 2018), we presented the sqRT-PCR data without statistically significance testing.

## RESULTS

To uncover a putative role for miR-184 in wound healing, we performed scratch assays on HPEK monolayers and examined miR-184 expression 5 days later, which was the time point at which we observed the highest expression of miR-184 in previous work (Richardson et al., 2018). As shown in Fig. 1a, a 50-fold induction of miR-184 was observed in scratched HPEK monolayers compared to their unscratched counterparts. We then determined whether SOCE contributed to the observed induction of miR-184. Incubation of scratched HPEK monolayers with SOCE inhibitors Gd^3+^ or BTP2 abolished the upregulation of miR-184 (Fig.1b,c). In contrast, when extracellular Ca^2+^ was raised to 1.5 mM to 5 days to evoke HPEK differentiation, only a relatively modest 10-fold rise in miR-184 expression was detected (Fig. 1d). Given that high Ca^2+^ induces miR-184 expression (Richardson et al., 2018), this suggests that further upregulation of miR-184 in differentiating HPEK is blunted compared to the scale of the response observed in proliferating cells. Nevertheless, it appears that miR-184 induction is a response to wounding in both proliferating and differentiating keratinocytes.

**Figure 1:**
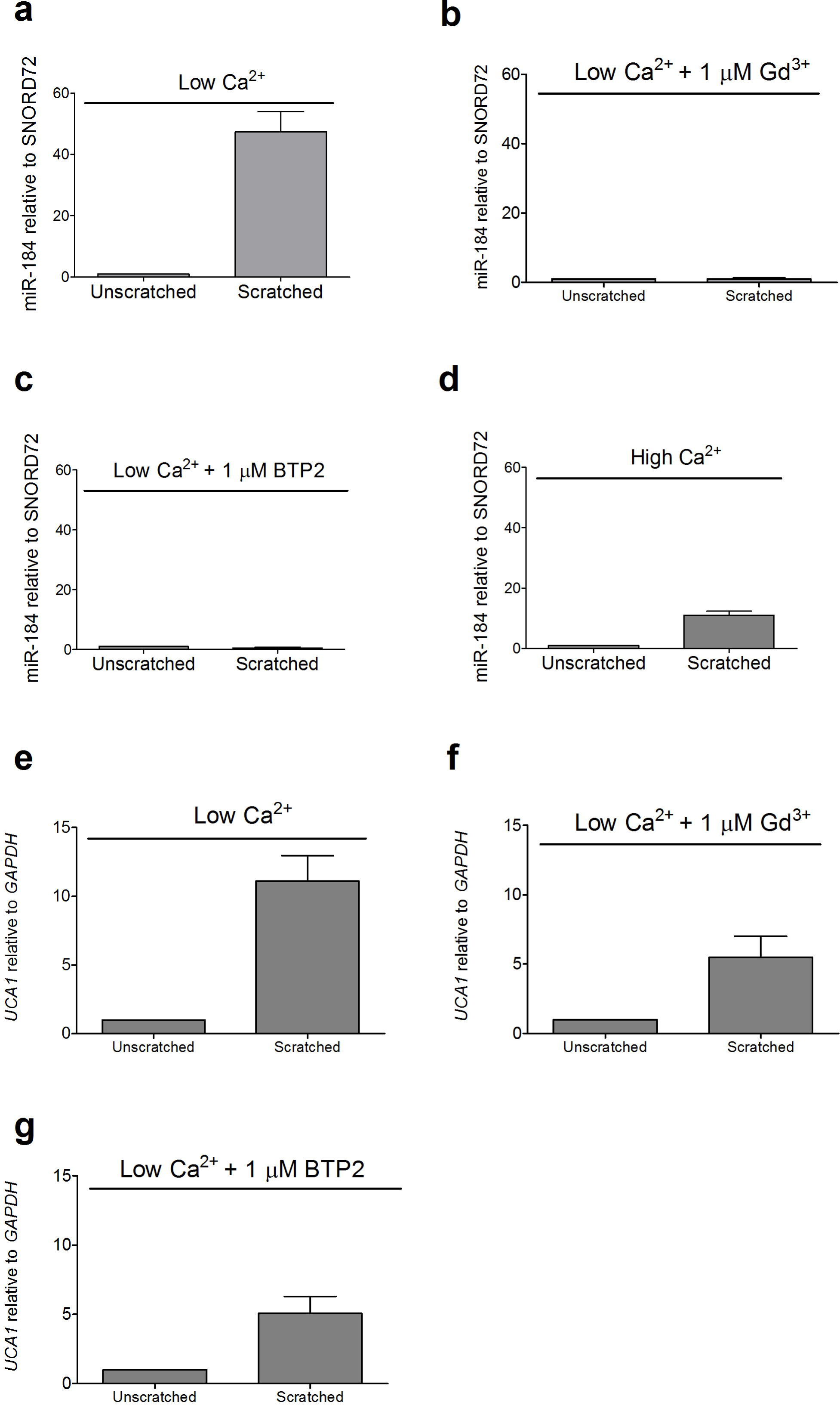
Induction of miR-184 in scratched monolayers. Confluent HPEK monolayers were scratched and cells harvested after 5 days in low (0.07 mM) or high (1.5 mM) Ca^2+^ media as indicated. The SOCE blockers Gd^3+^ and BTP2 were added 1 h before scratching and refreshed after day 2. (a-d) miR-184 expression normalized to SNORD72. (c-d) LncRNA UCA1 expression normalised to GAPDH. Data pooled from 3 independent experiments.

Given that UCA1 has been defined as a molecular sponge for miR-184, we considered the upregulation of miR-184 expression may be paralleled by an increase in UCA1. An 11-fold induction of UCA1 was observed in the scratched monolayers compared to their unscratched counterparts (Fig.1e). Further, blockade of SOCE with Gd^3+^ or BTP2 reduced UCA1 induction by around 50% (Fig. 1f,g). Thus UCA1 appears to be co-induced with its miR-184 target in wounded HPEK and the upregulation of UCA1 was at least partly dependent on SOCE.

We next evaluated the direct impact of miR-184 on HPEK migration. The cells were loaded with a miR-184 mimic and monolayers scratched as they approached confluence. Exogenous miR-184 led to a significant increase in the rate of HPEK migration compared to control cells loaded with a non-targeting oligonucleotide (Fig. 2a and Supplementary videos). In contrast, blockade of miR-184 activity using a locked nucleic acid (LNA) inhibitor impaired HPEK migration (Fig.2b). Interestingly, linear regression of the 12-36 h period from the relative migration rates (Fig. 2c) yielded migration indices of 0.53±0.11 per hour for miR-184-loaded HPEK, compared to 0.15±0.05, 0.13±0.03 and 0.04±0.02 per hour for cells loaded with control mimic, control inhibitor and miR-184 inhibitor, respectively. In other words, elevation of miR-184 appeared to enhance keratinocyte migration 3-fold while miR-184 inhibition decelerated keratinocyte migration 3-fold. Taken together, our results indicate for the first time that miR-184 is strongly upregulated in proliferating wounded keratinocyte monolayers and miR-184 activity regulates keratinocyte migration.

**Figure 2:**
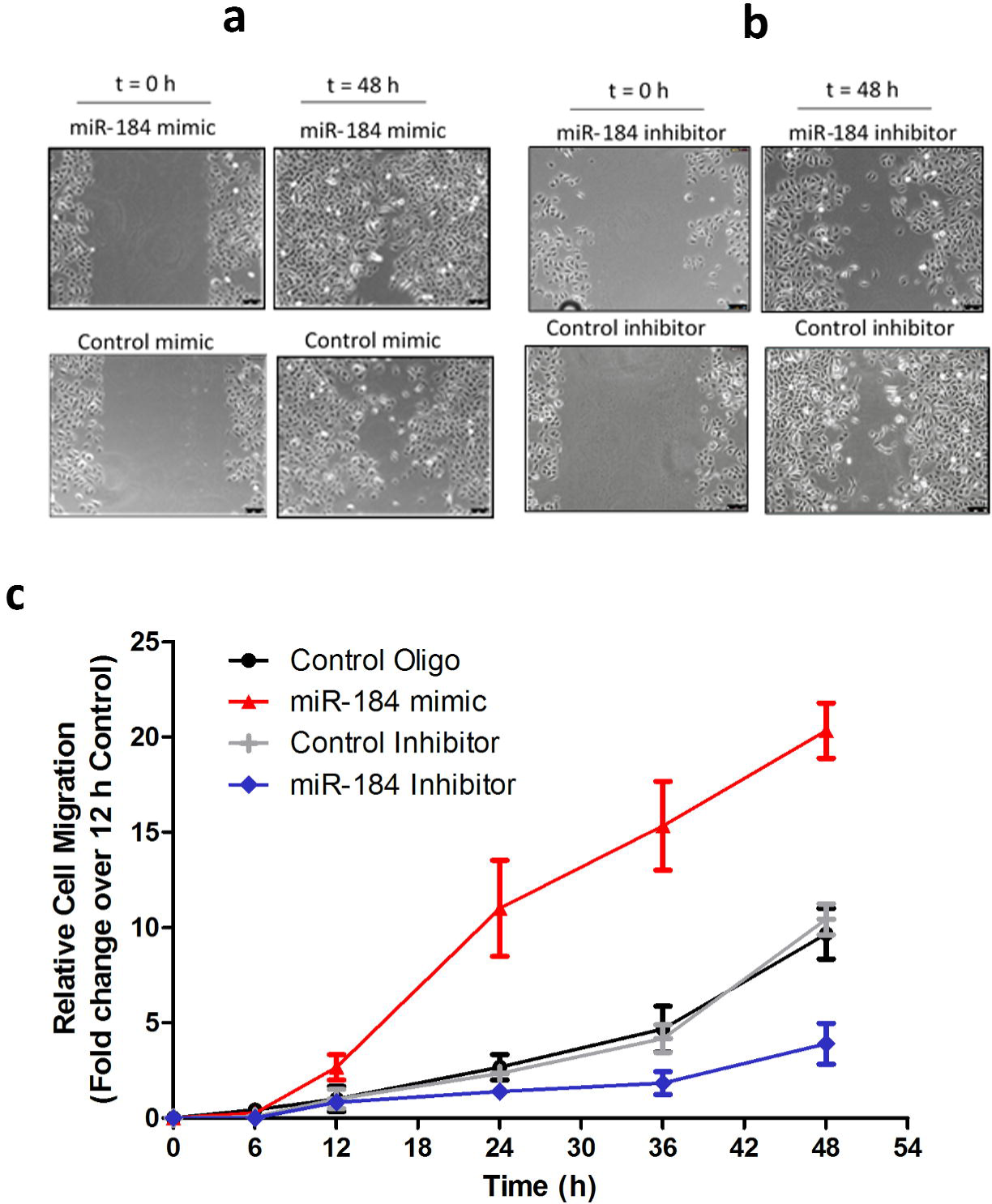
miR-184 promotes keratinocyte migration. Cells were loaded with 100 nM of miR-184 mimic (a), miR-184 inhibitor (b) or respective control oligonucleotides. Monolayers were scratched upon approaching confluence and monitored for 48 h. (c) The scratches were split into 3 equal sections and cells entering the middle were counted. Migration rates were normalised to the number of cells populating the breach at 12 h. Data pooled from 3 independent experiments.

## DISCUSSION

Non-healing wounds have persistently been associated with an increased risk of mortality (Boyko et al., 1996, Chammas et al., 2016, Walsh et al., 2016, Zarchi et al., 2015). Yet, only four therapies have received FDA-approval for chronic cutaneous wounds (Hamdan et al., 2017). Keratinocyte migration is crucial for the re-epithelisation of the wound and therefore molecular approaches that drive keratinocytes across a denuded wound surface have potential to enhance healing rates in patients. With miRNA-directed approaches gaining traction for the management of an increasing number of disorders (Rupaimoole and Slack, 2017), we investigated the involvement of miR-184 in keratinocyte migration.

We and others previously observed that miR-184 was largely absent from proliferating HPEKs under routine low Ca^2+^ culture conditions (Roberts et al., 2013, Yu et al., 2008). Consistent with this, recent work from the Shalom-Feuerstein laboratory indicated the miR-184 was low or absent in the basal layer of mouse epidermis, though the levels of miR-184 in cultured mouse and human keratinocytes were less clear (Nagosa et al., 2017). Nonetheless, our data suggest that wounding evokes *de novo* miR-184 expression that results in a 50-fold increase in steady-state miR-184 levels (Fig. 1). Mechanistically, SOCE appeared to be a key driver of the wound-dependent miR-184 expression, given that unrelated pharmacologic blockers of SOCE abolished miR-184 induction. However, the molecular signals that trigger SOCE following wounding of the HPEKs remain to be defined.

A growing number of studies have implicated UCA1 in the cell migration, mainly in relation to cancer but also, recently, in vascular smooth muscle cells (Tian et al., 2018). Given the proposed role of UCA1 as a ceRNA for miR-184 (Zhou et al., 2017), the concomitant induction of UCA1 and miR-184 in wounded HPEK monolayers suggests that UCA1 moderates the activity of miR-184 during migration. It will be interesting to determine whether modulation of UCA1 levels affects the migration of native and miR-184-loaded keratinocytes. Furthermore, as SOCE explains only part of the induction of UCA1, we await further experiments to fully define the axis from wounding to UCA1 induction.

From a translational perspective, the ability of exogenous miR-184 to accelerate HPEK migration suggesting miR-184 enhancement may promote re-epithelisation during wound healing. Recently, miR-132 was found to promote closure of diabetic mouse wounds and *ex vivo* human skin wounds (Li et al., 2017), showcasing the potential of miRNA-mediated wound healing. Questions remain though as to which vehicles are likely to support the safest and most efficient delivery of miRNA mimics to patient skin (Ross, 2018) and addressing these issues will catalyse the realisation of miRNA-based therapies to address unmet dermatological needs.

## ACKNOWLEDGEMENT

This study was funded by the British Skin Foundation (grant number 7006s).

## REFERENCES

Amelio I, Lena AM, Viticchie G, Shalom-Feuerstein R, Terrinoni A, Dinsdale D, et al. miR-24 triggers epidermal differentiation by controlling actin adhesion and cell migration. J Cell Biol 2012;199(2):347–63.

Bertero T, Gastaldi C, Bourget-Ponzio I, Imbert V, Loubat A, Selva E, et al. miR-483-3p controls proliferation in wounded epithelial cells. FASEB J 2011;25(9):3092–105.

Boyko EJ, Ahroni JH, Smith DG, Davignon D. Increased mortality associated with diabetic foot ulcer. Diabet Med 1996;13(11):967–72.

Chammas NK, Hill RL, Edmonds ME. Increased Mortality in Diabetic Foot Ulcer Patients: The Significance of Ulcer Type. J Diabetes Res 2016;2016:2879809.

Denzler R, Agarwal V, Stefano J, Bartel DP, Stoffel M. Assessing the ceRNA hypothesis with quantitative measurements of miRNA and target abundance. Mol Cell 2014;54(5):766–76.

Dykes IM, Emanueli C. Transcriptional and Post-transcriptional Gene Regulation by Long Non-coding RNA. Genomics Proteomics Bioinformatics 2017;15(3):177–86.

Eichhorn SW, Guo H, McGeary SE, Rodriguez-Mias RA, Shin C, Baek D, et al. mRNA Destabilization Is the Dominant Effect of Mammalian MicroRNAs by the Time Substantial Repression Ensues. Mol Cell 2014;56(1):104–15.

Eming SA, Martin P, Tomic-Canic M. Wound repair and regeneration: mechanisms, signaling, and translation. Science translational medicine 2014;6(265):265sr6.

Gudlur A, Hogan PG. The STIM-Orai Pathway: Orai, the Pore-Forming Subunit of the CRAC Channel. Adv Exp Med Biol 2017;993:39–57.

Guest JF, Ayoub N, McIlwraith T, Uchegbu I, Gerrish A, Weidlich D, et al. Health economic burden that wounds impose on the National Health Service in the UK. BMJ Open 2015;5(12):e009283.

Hamdan S, Pastar I, Drakulich S, Dikici E, Tomic-Canic M, Deo S, et al. Nanotechnology-Driven Therapeutic Interventions in Wound Healing: Potential Uses and Applications. ACS Cent Sci 2017;3(3):163–75.

Hogan PG, Rao A. Store-operated calcium entry: Mechanisms and modulation. Biochem Biophys Res Commun 2015;460(1):40–9.

Li D, Li X, Wang A, Meisgen F, Pivarcsi A, Sonkoly E, et al. MicroRNA-31 Promotes Skin Wound Healing by Enhancing Keratinocyte Proliferation and Migration. J Invest Dermatol 2015;135(6):1676–85.

Li X, Li D, Wang A, Chu T, Lohcharoenkal W, Zheng X, et al. MicroRNA-132 with Therapeutic Potential in Chronic Wounds. J Invest Dermatol 2017;137(12):2630–8.

Livak KJ, Schmittgen TD. Analysis of relative gene expression data using real-time quantitative PCR and the 2(-Delta Delta C(T)) Method. Methods 2001;25(4):402–8.

McShane BB, Gal D, Gelman A, Robert C, Tackett JL. Abandon Statistical Significance. https://arxivorg/abs/170907588 2017.

Nagosa S, Leesch F, Putin D, Bhattacharya S, Altshuler A, Serror L, et al. microRNA-184 Induces a Commitment Switch to Epidermal Differentiation. Stem cell reports 2017;9(6):1991–2004.

Nunan R, Harding KG, Martin P. Clinical challenges of chronic wounds: searching for an optimal animal model to recapitulate their complexity. Disease models & mechanisms 2014;7(11):1205–13.

Richardson A, Powell A, Sexton D, Parsons J, Reynolds NJ, Ross K. MicroRNA-184 is Induced by Store-Operated Calcium Entry and Regulates Early Keratinocyte Differentiation. https://doiorg/101101/319541 2018.

Roberts JC, Warren RB, Griffiths CE, Ross K. Expression of microRNA-184 in keratinocytes represses argonaute 2. J Cell Physiol 2013;228(12):2314–23.

Ross K. Towards topical microRNA-directed therapy for epidermal disorders. J Control Release 2018;269:136–47.

Rupaimoole R, Slack FJ. MicroRNA therapeutics: towards a new era for the management of cancer and other diseases. Nature reviews 2017;16(3):203–22.

Sen CK, Gordillo GM, Roy S, Kirsner R, Lambert L, Hunt TK, et al. Human skin wounds: a major and snowballing threat to public health and the economy. Wound repair and regeneration 2009;17(6):763–71.

Tian S, Yuan Y, Li Z, Gao M, Lu Y, Gao H. LncRNA UCA1 sponges miR-26a to regulate the migration and proliferation of vascular smooth muscle cells. Gene 2018.

Trafimow D, Amrhein V, Areshenkoff CN, Barrera-Causil CJ, Beh EJ, Bilgic YK, et al. Manipulating the Alpha Level Cannot Cure Significance Testing. Front Psychol 2018;9:699.

Usui ML, Mansbridge JN, Carter WG, Fujita M, Olerud JE. Keratinocyte migration, proliferation, and differentiation in chronic ulcers from patients with diabetes and normal wounds. J Histochem Cytochem 2008;56(7):687–96.

Vandenberghe M, Raphael M, Lehen’kyi V, Gordienko D, Hastie R, Oddos T, et al. ORAI1 calcium channel orchestrates skin homeostasis. Proc Natl Acad Sci U S A 2013;110(50):E4839–48.

Walsh JW, Hoffstad OJ, Sullivan MO, Margolis DJ. Association of diabetic foot ulcer and death in a population-based cohort from the United Kingdom. Diabet Med 2016;33(11):1493–8.

Wang T, Feng Y, Sun H, Zhang L, Hao L, Shi C, et al. miR-21 regulates skin wound healing by targeting multiple aspects of the healing process. Am J Pathol 2012;181(6):1911–20.

Wang T, Zhao N, Long S, Ge L, Wang A, Sun H, et al. Downregulation of miR-205 in migrating epithelial tongue facilitates skin wound re-epithelialization by derepressing ITGA5. Biochim Biophys Acta 2016;1862(8):1443–52.

Whittam AJ, Maan ZN, Duscher D, Wong VW, Barrera JA, Januszyk M, et al. Challenges and Opportunities in Drug Delivery for Wound Healing. Adv Wound Care (New Rochelle) 2016;5(2):79–88.

Yang C, Wu D, Gao L, Liu X, Jin Y, Wang D, et al. Competing endogenous RNA networks in human cancer: hypothesis, validation, and perspectives. Oncotarget 2016;7(12):13479–90.

Yang X, Wang J, Guo SL, Fan KJ, Li J, Wang YL, et al. miR-21 promotes keratinocyte migration and re-epithelialization during wound healing. Int J Biol Sci 2011;7(5):685–90.

Yu J, Peng H, Ruan Q, Fatima A, Getsios S, Lavker RM. MicroRNA-205 promotes keratinocyte migration via the lipid phosphatase SHIP2. FASEB J 2010;24(10):3950–9.

Yu J, Ryan DG, Getsios S, Oliveira-Fernandes M, Fatima A, Lavker RM. MicroRNA-184 antagonizes microRNA-205 to maintain SHIP2 levels in epithelia. Proc Natl Acad Sci U S A 2008;105(49):19300–5.

Zarchi K, Martinussen T, Jemec GB. Wound healing and all-cause mortality in 958 wound patients treated in home care. Wound repair and regeneration 2015;23(5):753–8.

Zhou Y, Wang X, Zhang J, He A, Wang YL, Han K, et al. Artesunate suppresses the viability and mobility of prostate cancer cells through UCA1, the sponge of miR-184. Oncotarget 2017;8(11):18260–70.

